# Fragment Linker Prediction Using Deep Encoder-Decoder Network for PROTAC Drug Design

**DOI:** 10.1101/2022.11.17.516992

**Authors:** Chien-Ting Kao, Chieh-Te Lin, Cheng-Li Chou, Chu-Chung Lin

**Affiliations:** AnHorn Medicines Co., Ltd., Taipei, 115202 Taiwan; Department of Biomedical Engineering, University of California Davis, Davis, CA 95616, USA

## Abstract

Drug discovery and development pipeline is a prolonged and complex process and remains challenging for both computational methods and medicinal chemists. Deep learning has shed light on various fields and achieved tremendous success in designing novel molecules in the pharmaceutical industry. We utilize state-of-the-art techniques to propose a deep neural network for rapid designing and generating meaningful drug-like Proteolysis-Targeting Chimeras (PROTACs) analogs. Our method, AIMLinker, takes the structural information from the corresponding fragments and generates linkers to incorporate them. In this model, we integrate filters for excluding non-druggable structures guided by protein-protein complexes while retaining molecules with potent chemical properties. The novel PROTACs subsequently pass through molecular docking, taking root-mean-square deviation (RMSD), the change of Gibbs free energy (Δ*G_binding_*), and relative Gibbs free energy (ΔΔ*G_binding_*) as the measurement criteria for testing the robustness and feasibility of the model. The generated novel PROTACs molecules possess similar structural information with superior binding affinity to the binding pockets in comparison to existing CRBN-dBET6-BRD4 ternary complexes. We demonstrate the effectiveness of AIMLinker having the power to design compounds for PROTACs molecules with better chemical properties.

## Introduction

Drug design is an iterative process with accounts of binding affinity, pharmacokinetics, and molecular structures to involve multiple cycles before optimizing a lead drug for trials. ^1^ Current challenge in structure-based drug design remains due to the size constraint of the search space and the logical drug-like molecules for chemical synthesis.^2^ Kick et al.^3^ demonstrate the ability to couple the complementary methods of combinatorial chemistry and structure-based design in a nanomolar range. Structure-based design also directs the discovery of a drug lead, which is not a drug product but, a compound with relatively superior nanomolar affinity for a target.^4^ Considering the current needs and limitations of drug discovery, the demand of expanding the knowledge into various targets is stretching.

Proteolysis-targeting chimeras (PROTACs) is one of the newly drawing attention modalities. PROTACs are hetero bifunctional small molecules, which connects a ligand for recruiting a target protein of interest (POI) and a ligand for an ubiquitin–protein ligase (E3), joint with an appropriate linker that further induces degradation of an target protein.^5,6^ Degradation is initiated when PROTACs promote the POI and E3 to form ternary complex.^7^ From a structural drug discovery point of view, PROTACs design relies on finding the best combinations of the three different chemical moieties and requires an attentive study of the structural characteristics of the E3 ligase and the POI complemented by molecular modeling and dynamic.^8,9^

Multiple E3 ubiquitin ligase have been targeted for PROTACs development and represents promising chemical properties in drug discovery. Here we focus on the CUL4-RBX1-DDB1-CRBN E3 ubiquitin ligase, which consists of Cullin-4 (CUL4), the RING-finger protein box1 (RBX1), the adapter Damage-specific DNA binding protein 1 (DDB1), and cereblon (CRBN) to form a macromolecular complex.^10^ Its substrate receptor cereblon (CRL4^CRBN^) binding to immunomodulatory drugs (IMiDs) may induce cancer therapeutical effect by targeting key neo-substrates to degrade.^11,12^ The principal of PROTACs recruiting E3 ubiquitin ligase and POI has been successfully applied to BRD4, a member of Bromodomain and Extra Terminal (BET) family that acknowledged in cancer for its role in organizing super-enhancers and regulating oncogenes expression.^13^ Winter et al. ^?^ designed the dBET6, a hybrid compound that drives the selective proteasomal degradation of BRD4 by linking to BET proteins and CRL4^CRBN^ ligand (hereafter called CRBN).^14^ The chemical properties have been proven to affect the structural rigidity, hydrophobicity, and solubility of PORTACs molecules.^15,16^ While studies have been made toward rational PROTACs design through structural biological and computational studies, linker design and generation still present a significant burden.

Current studies have leveraged the aid of rapid simulation and state-of-the-art deep learning for discovering novel structures, demonstrating the feasibility in timely and accurately screening the potential targets.^17,18^ Graph Neural Network (GNN) is one of the techniques gaining attention in drug discovery as it automatically learns task-specific representations by using graph convolutions and conserves the graph information as the atom-bond interactions.^19,20^ GNN learns the representations of each atom by aggregating the information from its surrounding atoms encoded by the atom feature vector, and recursively encode the connected bond feature vector through the message passing across the molecular graph, followed by a read-out operation that forms corresponding atoms and bonds.^21–23^ The state-of-the-art GNN models in predicting properties have been well demonstrated, and typically superior or comparable to traditional descriptor-based models. ^24,25^ In the study from Wu et al. ^26^ again showed evaluation results that GNN outperformed on most dataset, giving the network the feasibility of predicting various chemical endpoints. As such, GNN has been proven to be a potential model for designing and generating novel structures for drug discovery and investigating drug-like candidates.

Further, gated graph neural network (GGNN) performs best for molecular graph generation in deep generative models^27,28^ and demonstrates the practical structure generation in drug design.^19,29^ Many approaches use 2D SMILES-based chemical graphs embedded in low dimensional space, and generate new molecule by perturbing the hidden values of the sampled atoms.^30–33^ These studies are missing the nature of the molecular shape and the 3D information, which may greatly differ from the starting point of structure design. Another recent popular deep neural network drug design is in fragment linking technique. DeLinker, ^34^ adapted from Liu et al.,^31^ is the first attempt to apply GNN in linker design, particularly retaining the 3D structural information and generating linkers by giving two input fragments. 3DLinker^35^ puts the step forward with predicting the fragment nodes and sampling linker molecules simultaneously. However, none of these works have demonstrated an effective methodology of refining the generated molecule nor further considering validation in molecular conformations. The integration pipeline of adapting deep neural networks as the core technique in drug discovery and substantial validation process are still lacking in investigation.

In this work, we propose a novel deep learning based neural network, Artificial Intelligent Molecule Linker (AIMLinker). This network integrates designing, generating, and screening novel small molecular structures for PROTACs linkers, demonstrating a highly effective methodology of creating neo-structures to address the current difficulties in drug discovery. Our network considers the structural 3D information that first takes two fragments, with pre-defined anchors on both sides, and their structural information of angle and distance to represent the spatial positions between the input fragments. The core architecture of the network is GGNN^36^ with atoms and bonds denoting as nodes and edges, respectively. In addition, the iterative process of adding atoms and forming bonds is repeated until termination, followed by a read-out step for returning a newly generated compound and subsequently screening with the postprocess step. Outputs from the network are docked back to CRBN-BRD4 complex through AutoDock4 and validated with the measurement of the root-mean-square deviation (RMSD), the change of Gibbs free energy (Δ*G_binding_*), and the relative Gibbs free energy (ΔΔ*G_binding_*) to test the robustness and feasibility as a drug-like molecule. This end-to-end pipeline demonstrates a novel methodology for using state-of-the-art deep learning technique for drug discovery and shows the viability of designing novel PROTACs linker molecules.

## Methods

In this section, we first provide the details on preprocessing the POI and E3 ubiquitin ligase structures selected as the input for our encoder-decoder network. Next, we present the network architecture to generate the linker molecule with good viability and reasonable for drug synthesis. The postprocessing procedures are provided for validating the predicted molecules and conserving drug-like PROTACs molecules. Finally, the robustness of our predicted molecules is evaluated by docking and binding affinity metrics. The overall pipeline of our study is shown in Figure 1.

**Figure 1:**
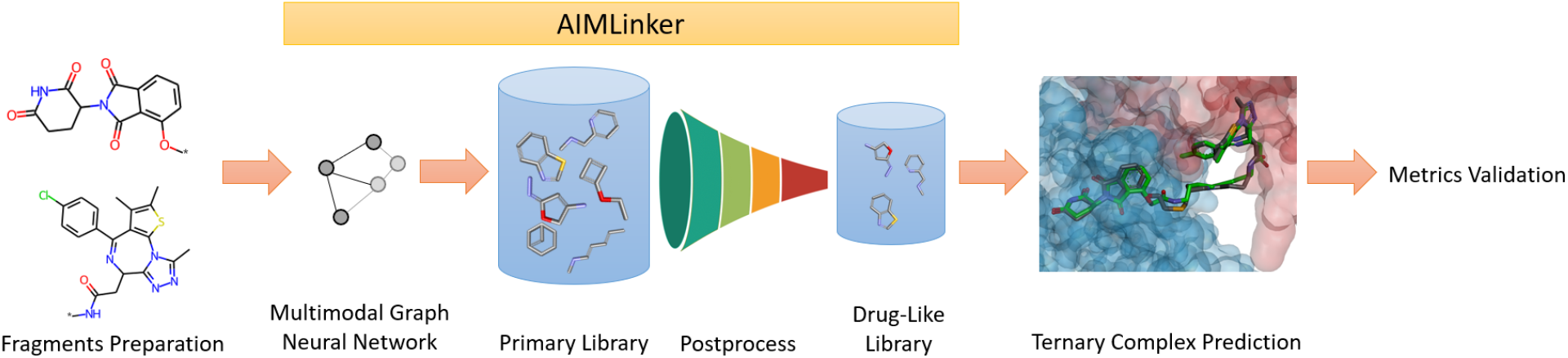
Scheme of the pipeline. The starting input data is two fragments, which are preprocessed and having their relative structural information. Next, AIMLinker takes the input fragments and generates the linker molecules. The molecules are postprocessed with our algorithm. We then take the best docking results of four molecules to validate their robustness of recognition as drug-like molecules.

### Data Processing

The plastic binding between the ligase and the substrate adopts distinct conformations depending on the linker length and position. Winter et al. ^?^ first showed dBET6, a PROTAC molecule, exhibiting high selectivity properties with the structure, and Nowak et al. ^37^ provided the ternary co-crystal structure of CRBN-dBET6-BRD4 (PDB: 6BOY) in Protein Data Bank. ^38^ The integration of structural, biochemical, and cellular properties of 6BOY ternary complex are designed to be a neo-degrader-mediated PROTAC structure. Figure 2A illustrates the relative spatial pose of the CRBN-dBET6-BRD4 ternary complex via Discovery Studio Visualizer (DSV)^39^ that red-labeled and blue-labeled structures are BRD4 and CRBN, respectively. dBET6 serves as the bridging molecule to link E3 ubiquitin ligase CRBN and target protein ligase BRD4.

**Figure 2:**
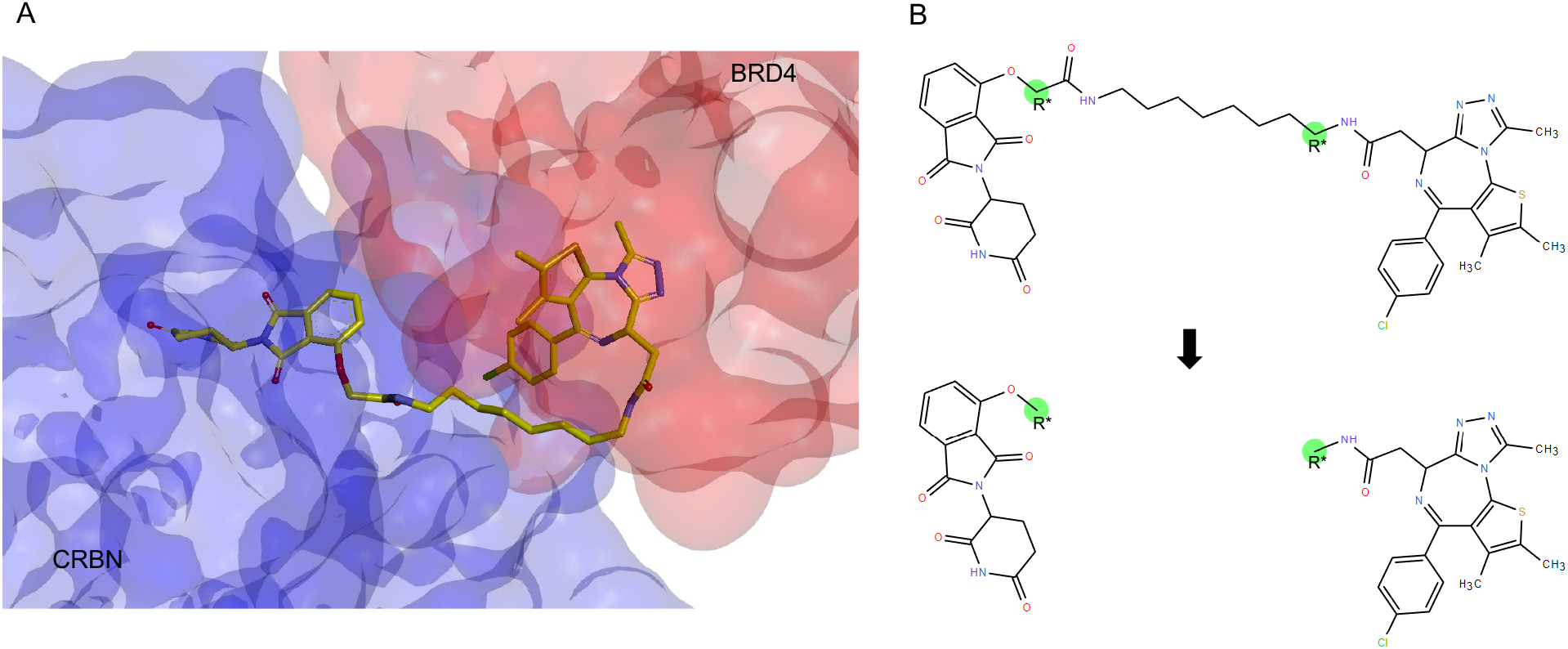
Scheme of the CRBN-dBET6-BRD4 ternary structure and processing protocol of dBET6. (A) The structure and the spatial information of 6BOY with the linker binding to BRD4 and CRBN.^37^ The red and blue label protein represent BRD4 and CRBN, respectively. (B) The 2D illustration of the dBET6 molecule links to E3 ubiquitin ligase and POI ligase. The anchors are highlighted and labeled with R* to feed the network with the start and end positions of the generated linkers. Next, the molecule between the anchors are removed and the remaining two fragments are considered as the input data for the network.

The input data for the network is two fragments composed of two ligands extracted from the PROTACs excluding the linker moiety. To prepare the input data from dBET6, we first retrieve the PROTACs molecule of the 6BOY structure. Considering the potent BRD4 inhibitor examined by Filippakopoulos et al.,^40^ the fragment of BRD4 ligand is defined as an illustration in Figure 2B, while the CRBN ligand is defined as a pomalidomide-like structure. The linking anchors on each ligand are labeled with R*, and the linker between these two anchors is removed. Since the co-crystal structure retrieved from PDB is spatially predefined and fixed, the anchors provide the 3D spatial information of the angle and distance between the two fragments. The network further takes the two fragments and the corresponding spatial information as the input to generate and design a linker library with the constraint of the space between the anchors.

### Multimodal Encoder-Decoder Network

We propose a novel network, AIMLinker, generating and designing novel structural linkers between fragments and postprocessing the predicted structures. The network is inspired by Imrie et al.^34^ and Liu et al.,^31^ taking two unlinked fragments, which contain their relative spatial position and orientation information, to generate the linker structures binding to the anchors on both fragments and form a novel molecular structure. This process is achieved by iteratively generating edges and adding new atoms from the selected pool, specifically 14 permitted atom types. The model generates the molecules in a breadth-first manner with a masking step to apply basic atomic valency rules. In addition, the network allows users to define the number of atoms between the anchors for maximizing the variations in generating the new linker molecules and provide the validity size of the two fragments corresponding to their distances. The other selection is the number of molecules to be generated and the network includes a postprocessing step for removing the molecules not subject to basic chemistry rules, duplicates, and illogical structures.

Figure 3 illustrates the iteration process, where the network uses a standard gated graph neural network (GGNN). The input fragments prepared from the data processing step are turned into a graph representation, where each atom and bond represent node and edge, and are labeled as *z* and *l*, respectively. A list of allowed 14 atom types is provided in the Supporting Information. The graph information is passed through AIMLinker, which utilizes GGNN as the core encoder structure, and the hidden state of nodes and edges are updated to integrate during the learning process (Figure 3a).

**Figure 3:**
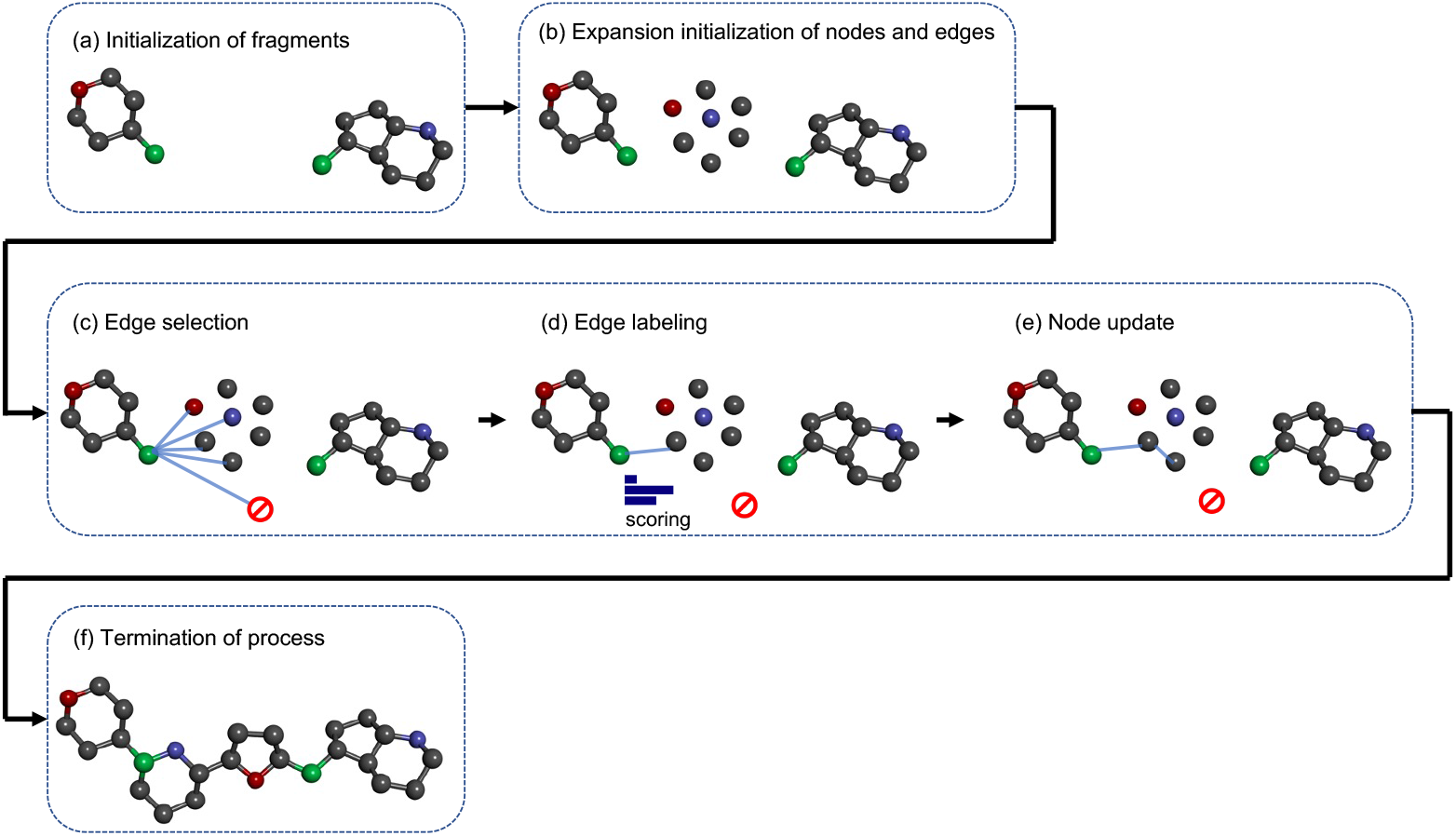
Network generation process. Two fragments in (a) are generated with the data processing steps, providing the spatial information, and the angle and distance between the anchors are calculated accordingly. Initialization of the nodes and edges in the network is illustrated in (b), where the 14 permitted atoms are randomly selected between the space. From (c) to (e) are the steps to process edge selection, edge labeling, and node updates, respectively. In particular, the three steps are sequentially repeated operations until the atom number reaches the maximum setting or all the edges and nodes are generated to cause (f) termination of the process.

Next, the graph representation of fragment input initializes with node expansion as shown in Figure 3b. Each node has a random hidden state *z_v_* derived from the *d*-dimensional normal distribution of 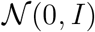, where *d* represents the number of features of the hidden state. The expansion nodes are subsequently labeled as *l_v_* association with structural information sampled from SoftMax output of a learned mapping *f*. The function *f* is applied with a linear classifier. However, it can be substituted with other functions to map the hidden state *z_v_* into different atom types *l_v_*. Note that the selected linker length can limit the number of expansion nodes.

New molecules are generated by iteratively selecting edges, labeling edges, and updating nodes as shown in Figure 3c-e. First, the initial node *v* considers whether to form an edge to the neighborhood node *u* in the graph. It is chosen by the start of the queue, which is configured by the initial input fragment anchors. The node is added to the queue if it is first connected to the graph. These processes are repeated until the expansion nodes are all queued, and no further nodes can be formed (i.e., termination of process in Figure 3f). The edge of the nodes *v* and *u* considers the basic valency constraint. The *f*, representing the core network GGNN, constructs a feature vector with the subsequent node *u* with such 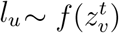. The edge between *v* and *u* at time point *t* is considered by a feature vector 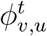:

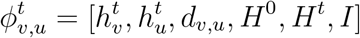

where 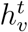 and 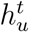 represent the hidden state of the initial node *v* and subsequent *u*, respectively. *d_v, u_* indicates the distance between the two nodes in the graph. *H*^0^ is the local information of the nodes, while *H*^t^ indicates the global information of the nodes at the time point *t*. *I* is encoded with 3D structural information of the relative angle and distance of the input fragments. The following representation is used to produce a distribution over the candidate edge:

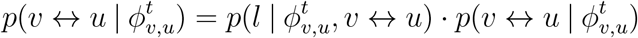

The edge between the two nodes *v* and *u* is formed in single, double or triple bonds after *u* is selected. Note that the bond formation is subject to basic valency constraints.

Lastly, all nodes are updated by their hidden state in accordance with the neural network in Figure 3e. We calculate the new hidden state 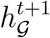 from initial hidden state 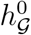 with considering the neighborhood nodes. At time point *t* + 1, we discard the previous hidden state 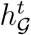, and conserve the local graph information of the queued nodes. This suggests that the molecule generation is independent of the graph history, and solely considers the local graph information. The iteration process will terminate when all the queues are empty. At the end of the generation, the largest intact molecule will be returned as shown in Figure 3f, and the unconnected nodes will be discarded. The stereochemistry information of the generated molecules is not given during the generative process. As such, a postprocessing step is needed for screening the predicted molecules and testing the robustness.

### Model Training

We prepare a dataset containing conventional ZINC dataset^41,42^ and PROTAC-DB,^43^ to train under variational autoencoder (VAE) framework. We construct the training dataset of 160,491 molecules with 157,221 and 3,270 from ZINC and PROTAC-DB, respectively. In the ZINC dataset, we select the chemical compounds with heavier and more complex structures. Meanwhile, in the PROTAC-DB dataset, all compounds to date are selected Each compound is segmented into two fragments and one linker as the input for the network. We construct the linker segment to contain at least three heavy atoms while retaining the intact ring structures in the linker or the two fragments. Next, we split the dataset into 90% for training and 10% for the validation process by adapting 10-fold cross-validation to overcome the overfitting issue. We tune the hyperparameter as reflected in Table S1 to retrieve the best performance.

The model trains on the dataset focusing on the fragment-molecule pairs. Given the two input fragments A and linked molecule B, the goal of the model is to reconstruct B from (A, *z*).The original linked molecule B is transferred into graph representation and *z* is the latent code of the learned mapping. Specifically, z is learned from a set of expansion nodes of B, and the mapping is averaged into a low-dimensional vector. We constrain *z* into the low-dimensional vector to enforce the network learning the information from B and follow to degenerate the network for B. The loss function is similar to standard VAE loss including a reconstruction term on a Kullback-Leibler regularization, and a regression loss: 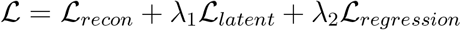. Note that we allow variations from the pure VAE loss (*λ*_1_ = 1) by Yeung et al.^44^

### Postprocessing

The raw outputs from the model are PROTACs library in 2D chemical structure and are further postprocessed with our screening procedures to remove the unfavored targets. Figure 4 shows the filters of our proposed method that are integrated into the AIMLinker. Due to the constraint of the graph computational process and the linked substructures, the model generates the molecules by choice, which includes duplicated predictions, undruggable targets, and structures violating the basic chemistry law. The first filter rules out the duplicates, the same structures predicted by the model, leaving every unique molecule after this process. This process also includes the non-linker substructures, i.e., two fragments are not formed into one compound with the linker moiety. Next, we have a library with unfavorable substructures that are not feasible for chemical synthesis, or unable to become a druggable target. This library includes the substructures such as acid halide, disulfide bond, peroxide bond, and small-number cyclic rings with double bonds, and additionally by the model to predict whether the newly generated substructures are feasible as a drug lead. This step is significantly important for screening the target pool to reduce the molecule numbers while retaining the candidates potentially having high binding affinity and chemical activity. Finally, the molecules that violate Bredt’s Rule,^45^ which the substructures contain certain bicycle atomic-bridged-ring structures with a carbon-carbon double bond at a bridgehead atom, are removed from the target pool. These steps remove the unwanted molecules from the target pool and the remaining targets remarkably reduce the needed computational resources and the time span for simulation.

**Figure 4:**
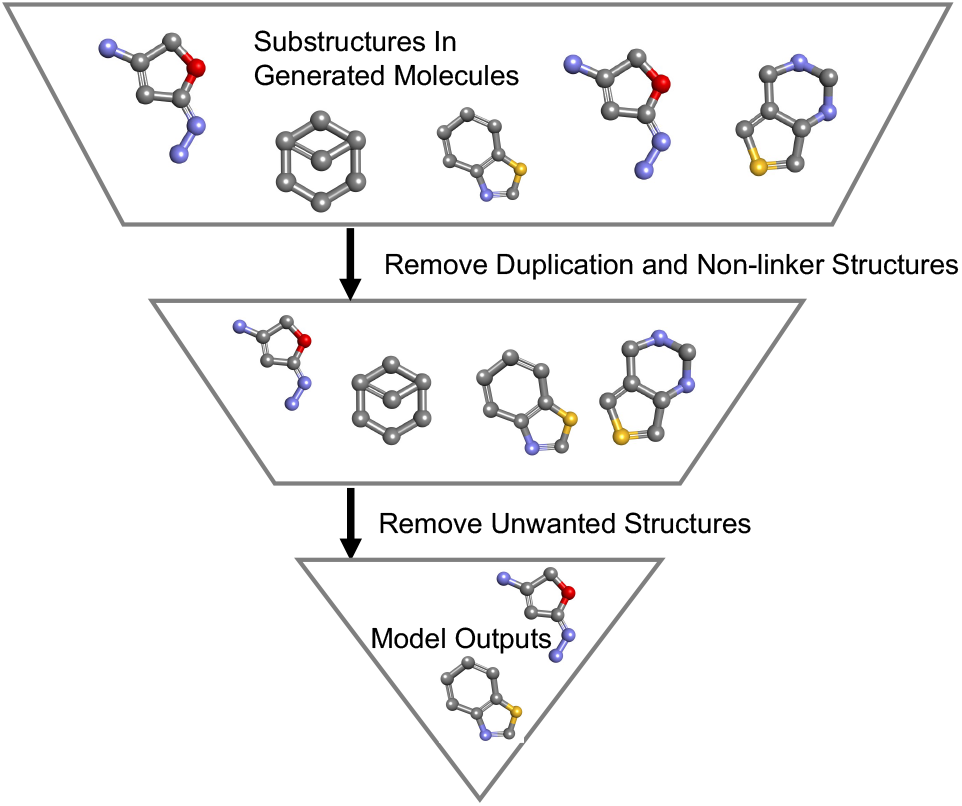
Workflow of postprocess. The generated molecules pass through multiple steps of filters, specifically, removing duplicates, non-linker, and unwanted substructures. This postprocess step is integrated into AIMLinker.

To further validate the merit of using postprocess steps, we utilize the “Rule of Three” to measure the effectiveness of our generated molecules. “Rule of Three” refers to the molecular weight (MW) of a fragment being <300 Dalton, the calculated logarithm of the 1-octanol–water partition coefficient of the non-ionized molecule (cLogP) being ≤ 3, the number of hydrogen bond donors (HBD) being ≤ 3, and the number of hydrogen bond acceptors (HBA) being ≤ 3, and we also include the polar surface area (PSA) being ≤ 60 Å in addition to the standard setting of the rule.^46,47^ We apply this rule to the generated linker pool to measure and calculate the molecular properties at each filter step.

### Docking Validations

Before applying docking methods, 3D protein-protein interaction poses and 3D conformations of postprocessed molecules need to be constructed first. The co-crystal structure of CRBN-dBET6-BRD4 is released in PDB and can retrieve the simulated spatial conformation via DSV. Meanwhile, all postprocessed molecules initially sketched as 2D chemical structures, are converted into 3D PROTACs conformations through DSV. The reference compound dBET6 is also reconstructed into a series of 3D conformations for validating the consistency of our docking methodology. These 3D compounds are subsequently minimized using the energy minimization method.^48^ After protein-protein interaction poses and 3D conformations of PROTACs candidates are well prepared, AutoDock4^49^ is applied to predict the best PROTACs binding pose by labeling the binding pocket as a grid. During the docking procedure, each 3D PROTACs freely binds to CRBN and BRD4 with consideration of the binding energy, biochemistry property, and entropy. We allow 10 binding poses of each PROTACs from the network and form a 10-time dataset as the initial docking inputs. As such, our PROTACs library and dBET6 can freely rotate, fold, and bind to the pocket to form the best pose in DSV with the highest binding affinity and lowest entropy energy.

To validate the robustness of generated molecules from AIMLinker and dBET6 provided in PDB, we measure the metrics including structural information and binding affinity. We use RMSD, which was introduced by Bell et al. ^50^ for measuring between respective atoms in two molecules,

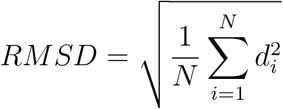

where *N* is the number of atoms in the ligand, and *d_i_* is the Euclidean distance between the *i^th^* pair of corresponding atoms. We take the crystal structure in PDB as the reference, and measure the structural similarity level with the generated linker molecules by superimposing and calculating the RMSD values.

Another metric taken into consideration for validating the molecules is Δ*G_binding_* of the protein–ligand complex. It is calculated from molecular mechanics Poisson–Boltzmann surface area (MM-PBSA) method.^51^ The MM-PBSA approach is one of the most widely used methods to compute interaction energies among biomolecular complexes. In general, Δ*G_binding_* between a protein and a ligand in solvent can be expressed as:

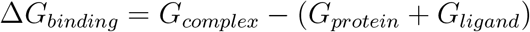

where *G_complex_* is the total free energy of the protein-ligand complex and *G_protein_*, and *G_ligand_* are the total free energies of the separated protein and ligand in the solvent, respectively. We individually generate 10, 25, 50, and 100 poses of each molecule, and AutoDock4 determines the best logical spatial orientation of each trial independently. The best pose of each trail is retrieved and applied to the docking process to calculate the Δ*G_binding_*. The metric is averaged between trials with variations. Note that we constrain the CRBN-BRD4 spatial position and consider the free movement of the generated bridging molecules.

### Molecular Dynamics Simulation

To further examine the robustness of the generated molecule, we take the best linker structure to compare its Δ*G_binding_* and ΔΔ*G_binding_* with redocked dBET6. The binding affinities of the three selected PROTACs (dBET6 crystal pose, dBET6 redocked pose, and 6boy_1268) with CRBN-BRD4 are further evaluated with 10 ns MD simulations by GROningen MAchine for Chemical Simulations (GROMACS) 2022.4 version.^52^ Each CRBN-PROTACs-BRD4 ternary complex was constrained using the CHARMm force field^53^ and solvated using the transferable intermolecular potential with 3 points (TIP3P) water molecules in a truncated octahedron water box with a minimal distance of at least 10 Å from any edge of the box to any protein atom. Each CRBN-PROTACs-BRD4 complex was initially energy minimized to remove the unfavorable contacts using the conjugate gradient method for 10,000 steps, and subjected to 100 ps of heating from 50 K to 300 K. Subsequently, a 500 ps equilibrium run was performed. Finally, periodic boundary dynamics simulations of 10 ns were conducted for the production step in a NPT ensemble at 1 atm and 300K. During the simulations, all covalent bonds involving hydrogen were constrained using the SHAKE algorithm. The long-range electrostatic interactions were treated using the particle mesh Ewald method. ^54^ The output trajectory files were saved every 2 ps from a 10 ns period and used for subsequent binding free energy analysis.

To calculate binding free energy Δ*G_binding_*, the 5,000 frames from 10 ns trajectory of CRBN-dBET6 (crystal pose)-BRD4, CRBN-dBET6 (redocked pose)-BRD4, and CRBN-6boy_1268-BRD4 ternary complex are individually calculated and averaged by applying the MM-PBSA method. The relative binding free energy ΔΔ*G_binding_* is measured by comparing the dBET6 crystal pose with redocked dBTE6 and 6boy_1268, respectively.

## Results and Discussions

We develop a novel network, AIMLinker, generating neo-structure of small molecule linker for PROTACs degradation protein. AIMLinker takes two fragments with structural information as the input data and processes with a deep learning network to create linker molecules. We first provide the details of the generated molecules with their chemical properties and structural statistics. Next, we take four molecules with the highest binding affinity to compare dBET6 with their RMSD and ΔG.

AIMLinker demonstrates a robust and rapid pipeline for generating and designing new PROTAC linkers. Our network combines two processes into one end-to-end pipeline: 1) takes two unlinked fragments as input and uses an encoder-decoder deep learning network to generate the substructures forming a new PROTACs molecule. We consider the structural information in the form of the angle and distance between the two initial fragments, and iteratively add atoms between the space until filled or reach the limitation of maximum atom setting, 2) postprocesses the generated molecules to extract the potential drug-like molecules. In particular, we screen with duplicates, and exclude structures violating basic chemistry rules and unwanted substructures. This rapid pipeline provides the viability of timely generating novel small molecules with high binding affinity to CRBN-BRD4, and the potential to translate the work to other PROTACs targets.

### Generated Molecules

We generate the molecules with a set of range between the fragments. This range gives flexibility to the network in designing a more linear or ring-like structure. The raw output from the neural network is 2,000 structures, including illogical molecules and unwanted substructures. Therefore, we then take these outputs to undergo postprocess procedures with two filters applied: 1) first filter removes the duplicated molecules and non-linker structure, i.e., two fragments are not formed into one compound with the linker structure. After this filter, the remaining number of molecules is 1,175. The remaining molecules in this filter are unique and novel, yet not favorable to become a drug lead, 2) filter out the unwanted substructures that are not applicable to drug-like molecules. The final output number from AIMLinker is 524 molecules.Note that our model generates new and effective molecules with a comparable successful rate to other state-of-the-art ML methods in Table S1. We highlight the second postprocess screening approach to retain the druggable and potential drug leads.

Next, we utilize the “Rule of Three” to validate the effectiveness of applying postprocess steps in the linker structure. Table 1 shows the rule of three metrics of MW, cLogP, HBD, and HBA, and we additionally include PSA here. The generated pool of molecules applied with postprocess step outperformed that of preprocessing step except cLogP. Specifically, our proposed method has 93%, 95%, 95%, 60%, and 48% of the molecules that pass the rules in MW, cLogP, HBD, HBA, and PSA, respectively. For preprocessed molecular pool, it achieves 91%, 97%, 90%, 49%, and 36% in the corresponding metrics. In addition, this method surpasses the preprocessed data with as high as 12% in PSA, while the lowest surpassed percentage compared to the preprocessed data is 2% in MW. We show better performance with this additional postprocess step in 4 out of 5 metrics, demonstrating the robustness of the linker molecules possessing better chemical properties.

**Table 1:** Results of “Rule of Three” parameters analysis. Abbreviations: molecular weight, MW; the calculated logarithm of the 1-octanol–water partition coefficient of the non-ionized molecule, cLogP; number of hydrogen bond donors, HBD; number of hydrogen bond acceptors, HBA; polar surface area, PSA.

| Parameters | Preprocess (%) | Postprocess (%) |
| --- | --- | --- |
| MW ($<300$ Da) | 91 | **93** |
| cLogP ($\leq 3$) | 97 | 95 |
| HBD ($\leq 3$) | 90 | **95** |
| HBA ($\leq 3$) | 49 | **60** |
| PSA ($\leq 60\text{\AA}^2$) | 36 | **48** |

Table 2 provides the structural statistics of the final output from AIMLinker. In particular, the generated molecules from AIMLinker provide ring-shape structures, while dBET6 linker is a linear structure, which gives the compound more flexibility to freely rotate inside the binding pockets and the opportunities of binding to other positions to reduce the compound potency and the pharmacokinetics property. Our generated linker structures between the two input fragments provide 229 ring-like substructures out of 524 molecules, 43% of the total number. Of the 229 compounds, there are 32 have bicyclic rings, and one compound has tricyclic rings. The incidences of ring-shape structures of the designed molecules are demonstrated in Table 2 that the numbers of three-membered, four-membered, five-membered, and six-membered rings, and the number of atoms in the ring structure above 6 are 24, 30, 90, 112, and 6, respectively. These cyclic compounds restrict the rotational angles and the possibility of binding to non-target binding positions. In addition, the ring-link structures generated by AIMLinker provide more stability for the compound and possess the ability to form strong π bonds increasing the binding affinity in the binding pockets.

**Table 2:** Ring-structure statistics of the generated molecules. Here we provide the number of incidences of different numbers of membered rings.

| Ring Structures | Number of Molecules |
| --- | --- |
| 3-membered Ring | 24 |
| 4-membered Ring | 30 |
| 5-membered Ring | 90 |
| 6-membered Ring | 112 |
| Above 6-membered Ring | 6 |

**Table 2:** Docking performance of redocked dBET6 and the generated molecules. We individually generate 10, 25,, 50 and 100 poses of each molecule, and select the best pose via AutoDock4 simulation. The metrics are shown in average of the four trials with variations.

| Linker Molecules | Average RMSD (Å) | Average $\Delta G_{binding}$ (kcal/mol) |
| --- | --- | --- |
| dBET6 (crystal pose) | — | −22.51 |
| dBET6 (redocked pose) | $0.59 \pm 0.01$ | $-54.67 \pm 11.53$ |
| 6BOY_1268 | **$0.44 \pm 0.05$** | **$-62.26 \pm 6.36$** |
| 6BOY_1974 | $0.46 \pm 0.06$ | $-55.33 \pm 4.23$ |
| 6BOY_1854 | $0.58 \pm 0.07$ | $-56.86 \pm 8.07$ |
| 6BOY_0518 | $0.77 \pm 0.08$ | $-50.41 \pm 5.23$ |

### Docking Performance

To assess the generated molecules and compare them with the existing dBET6 structure, we use AutoDock4 for docking and validation. In order to consolidate AutoDock4 having the ability to be a reference tool for measuring the generated molecules, we redock the compound to the binding pockets of CRBN and BRD4. We constrain the free energy of dBET6 to retrieve the closest docking pose and binding affinity provided in the 6BOY crystal structure. The final output of 524 molecules from AIMLinker are then passed through AutoDock4. In the standard protocol and matching with the biological interaction, we allow a maximum of 10 binding poses for each molecule. Since not every molecule is feasible to bind within the pocket, the total is 5,095 poses generated from docking. We set the RMSD threshold value of less than 1 Å to be considered as drug-like molecules. The four displayed molecules in Figure 5 are extracted from this threshold.

**Figure 5:**
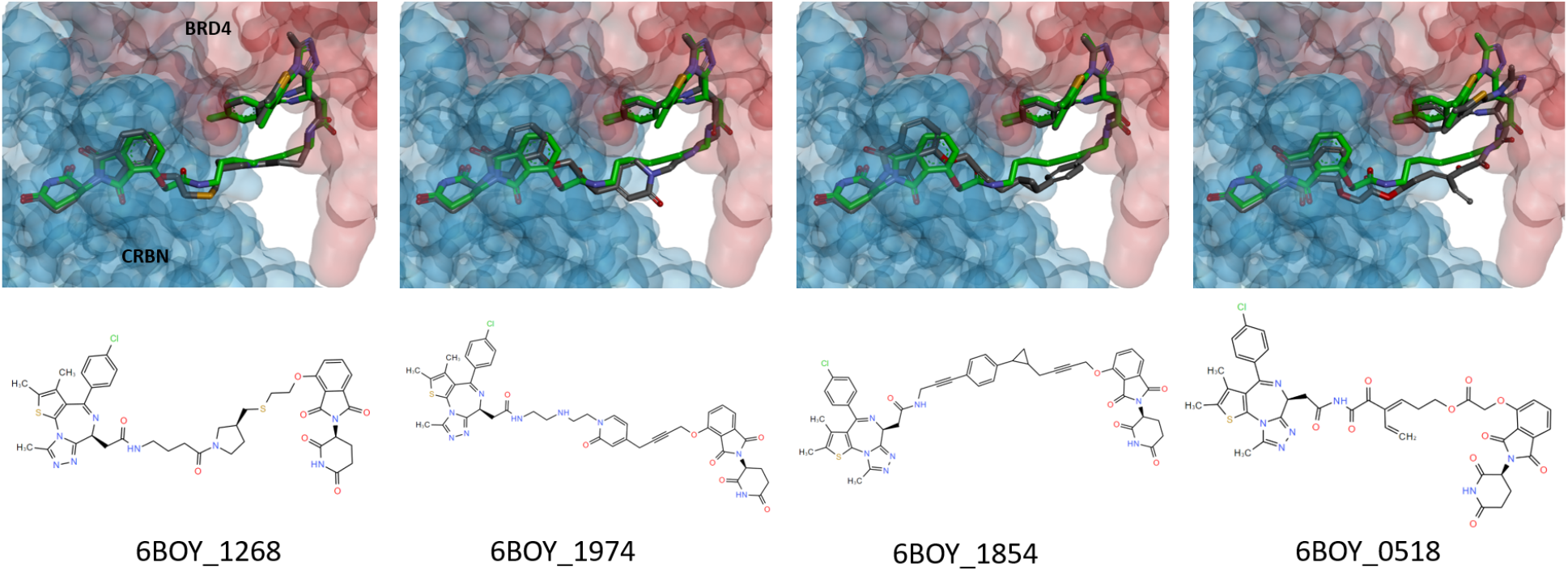
Docking poses of the generated molecules that the RMSD value less than 1 Å, which takes dBET6 conformation as the reference compound.

In Table 2, we show the average RMSD and average Δ*G_binding_* values. The spatial structural information is measured in RMSD values. Of the re-docked dBET6 and four generated molecules, 6BOY_1268 has the best average RMSD of 0.44 Å, and 6BOY_1974 performs the second best of 0.46 Å. For the average Δ*G_binding_*, both 6BOY_1268 and 6BOY_1974 possess better binding affinity. The performance suggests that the generated molecules have superior chemical properties than either the crystal pose or redocked pose of dBET6 molecule. Further, the third-best molecule 6BOY_1854 also exhibits comparable chemical properties to the redocked dBET6. The results demonstrate that the model generates novel and comparable molecules in PROTACs drug design.

### Molecular Dynamics Simulation

After molecular docking, the CRBN-dBET6 (crystal pose)-BRD4, CRBN-dBET6 (redocked pose)-BRD4, and CRBN-6boy_1268-BRD4 ternary complexes were subjected to 10 ns MD simulations, and the binding free energies were calculated, as reported in Table 3. From the average Δ*G_binding_* value, the complexes formed between 6boy_1268 and CRBN-BRD4 present the lowest calculated values, suggesting that 6boy_1268 forms the most stable complexes with CRBN-BRD4 in compared to the values of the benchmark compound (dBET6 crystal pose and redocked pose). Further, the ΔΔ*G_binding_* of 6BOY_1268 is lower than redocked dBET6 pose to indicate a better binding affinity in CRBN-BRD4 pocket. These results further consolidate the robustness of 6BOY_1268 possessing better chemical properties than redocked pose, and potentially becoming a potent drug target.

**Table 3:**
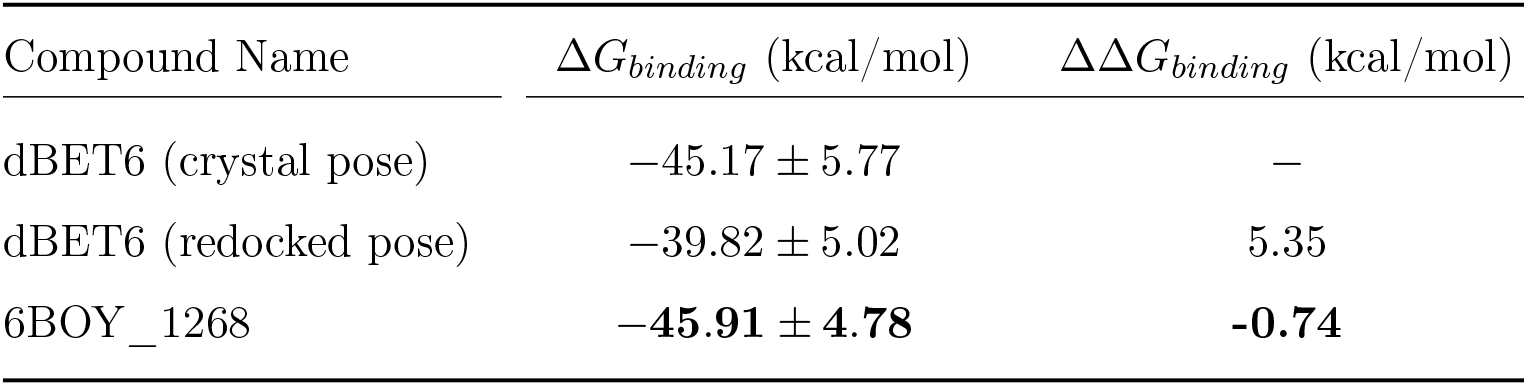
Average Δ*G_binding_* and ΔΔ*G_binding_* values for the crystal structure dBET6, redocked dBET6, and generated linker with best chemical properties.

## Conclusion

In this work, we propose a deep neural network to generate and design novel PROTACs molecules. We integrate sampling and postprocessing steps to extract the potent druglike molecules and demonstrate the robustness of the generated molecules. We show that our generated structures possess superior chemical properties to the existing compound. Our model can perform virtual high-throughput screening for rapid generation and reducing manual labors. We focus on a single PROTACs target for testing and validating our proposed model. In the next stage, we aim to expand the utilization of the model and apply it to more PROTACs targets, and further investigate the applications in other fields of drug discovery.

## Data Availability Statement

All data mentioned in this study are publicly available at ZINC dataset, PROTAC-DB, and PDB. We retrieved the training and validation data from above databanks. All the data we applied can be found at the supplementary document and https://github.com/AnHorn/AIMLinker.

## Supporting Information

Details on the neural network implementation, benchmark comparison to other methods, and data preparation. AIMLinker hyperparameter tuning in Table S1; performance on filtering generated molecules by using other ML methods in Table S2; structures of postprocess filters in Figure S1; truncated side chain of 3DLinker input fragments in Figure S2.

---

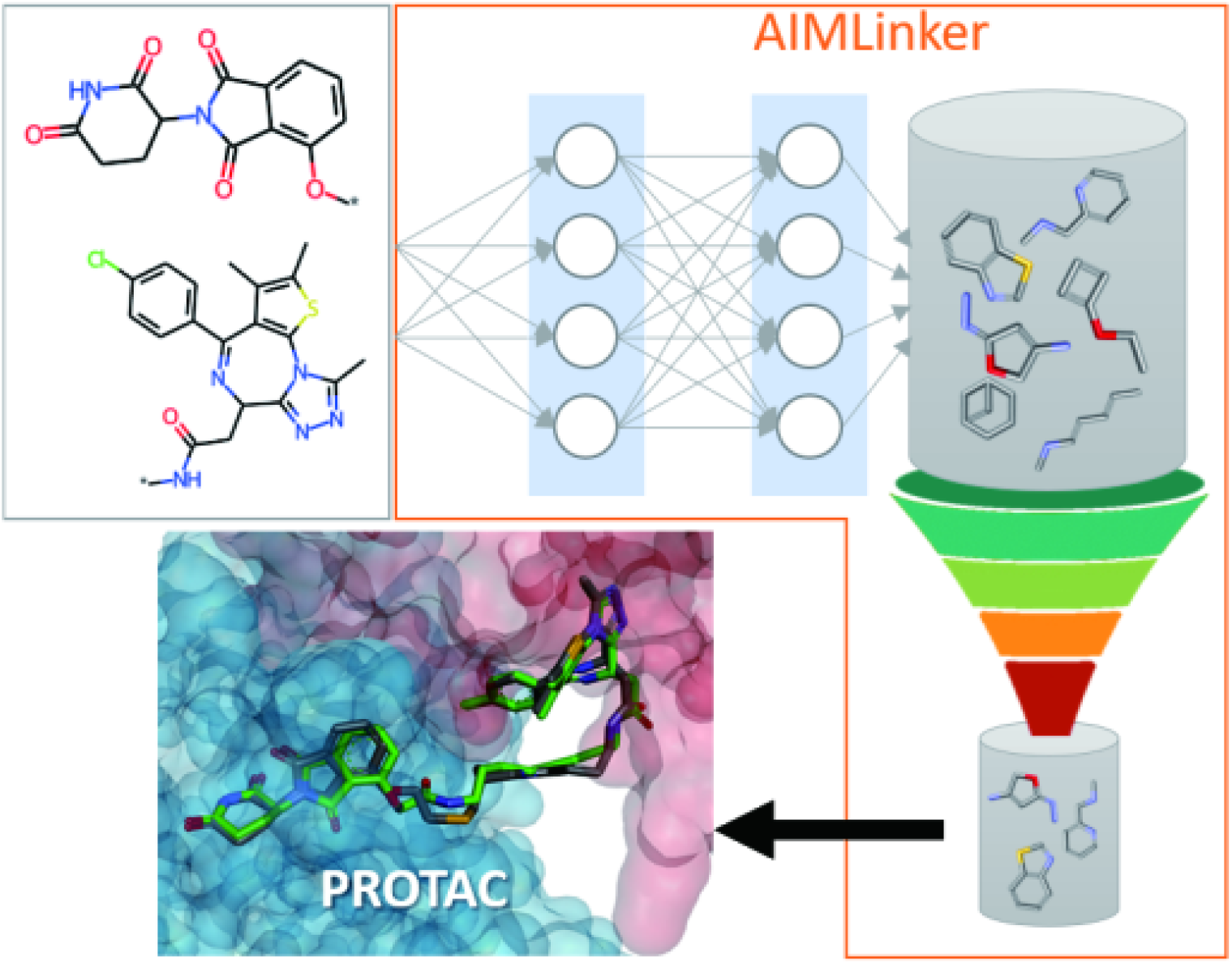

