## Supplementary Material for "Fragment Linker Prediction Using Deep Encoder-Decoder Network for PROTAC Drug Design"

### Data Preparation

We remove the linker of dBET6 through Discovery Studio Visualizer (DSV) to generate two fragments as input for the neural network. The position of each ligand is determined as anchors and labeled with $R^*$. Next, the distance and angle between the anchors are calculated and fed into the network as a feature vector. In addition, we construct the training dataset using the same method by removing the linker and determining the fragments. The goal of the learning process from the input is to reconstruct the intact molecule from the input fragments.

### Multimodal Encoder-Decoder Network

#### Training Set

PROTAC-DB,[1] a public web-based database collecting chemical structures and biological activities of PROTACs, contains 3,270 PROTACs to date. All structures are downloaded and converted into SMILES as the training set. On the other hand, our ZINC dataset is similar to that selected by Imrie et al.,[2] was also taken into consideration to enrich structural diversity. With that, we constructed a total of 160,491 structures as the training set.

#### Permitted Atoms

During the optimization and generation, the nodes are labeled with $l$ to indicate the atom types. There are 14 atom types allowed to update in the process, including carbon, nitrogen ($N^-$, N, $N^+$), oxygen ($O^-$, O, $O^+$), sulfur (maximum valence are 2, 4, 6), fluorine, chlorine, bromine, and iodine.

#### Network architecture

In our network, we use a standard gated graph neural network (GGNN) as the backbone training with public ZINC and PROTAC-DB to tune the model. We adopt L1 and L2 regularization to overcome overfitting issues during training. Furthermore, Adam optimizer is used in the process. For training and optimizing the model weights, the two datasets combined with around 160,000 molecules, so we do not apply an augmentation step for the training process. We also monitor the training and validation loss to ensure the model continuously learns useful features for further generating novel molecules.

#### Hyperparameter settings

We perform hyperparameter tuning to retrieve the best performance. Since the neural network is generating similar performance throughout the trials, we tune with limited hyperparameters. The details are as follows:

Table S1: Hyperparameter tunning. The bold texts are the values we selected for the best validation performance.

| Parameters | Settings |
|---|---|
| Learning rate | 0.1, 0.01, **0.001**, 0.0001 |
| Batch size | 8, **16**, 24, 32 |
| Hidden state dimension | 16, **32**, 50 |
| Encoding dimension | 2, **4**, 8 |

### Postprocessing Filters

Postprocessing, integrated with RDKit,[3] is composed of multiple steps of filters to remove duplication, non-linker, and unwanted substructures. Molecules with unfavorable substructures, containing but not limiting as Figure S1 shows, are removed. Our postprocessing script is released on our GitHub repository.

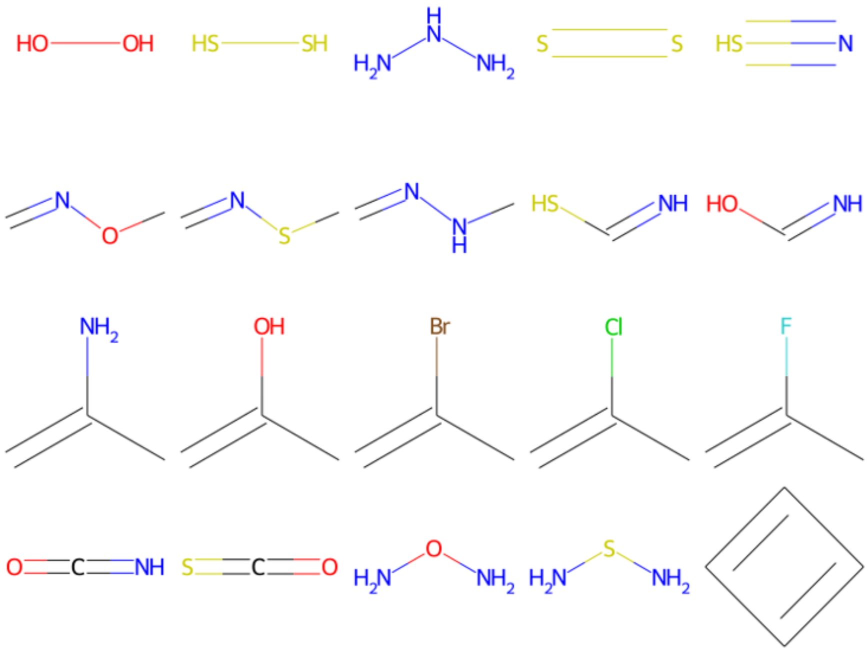


Figure S1: Scheme of unfavorable substructures.

### Benchmark

We compare AIMLinker with other state-of-the-art ML methods, including DeLinker,[2] 3DLinker,[4] and DiffLinker.[5] We use the same procedure in AIMLinker to generate the input fragments at the data preparation step. Next, apply the postprocess filters to retrieve the potential drug-like molecules. Table S2 shows the assessment of the number of outputs from each network, the number of outputs after removing duplicates, and the number of outputs after removing unlinked structures. Further, more than half the number of generated structures from DeLinker are unlinked to the anchors. This might result from the length of the linker in PROTACs being much longer than the general small molecules trained in DeLinker.

3DLinker (Github commit number: b8f5c92) fails to apply the same fragment with anchors as we used in AIMLinker. During the preparation step, 3DLinker takes SMILES as input and generates a minimized conformation; however, the energy minimization of the dBET6 conformation is not converged so no conformation is generated. We thus used side-

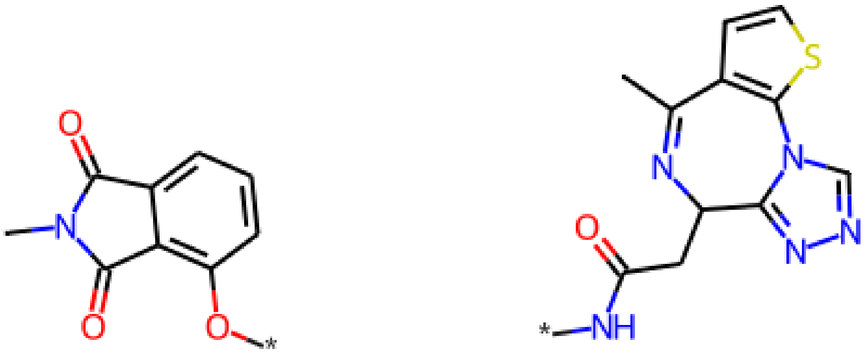

Figure S2: Truncated side chain used in 3DLinker input.

chain-truncated fragments as shown in Figure S2 to enable 3DLinker functionality. The linker length cannot be set in 3DLinker, so we calculate the linker length on the output with a major number of linker lengths of 9-11 heavy atoms (not including both anchors). A shorter length improves the unlinking issue, but the PROTACs with insufficient length reduce the variations to the generated molecules. Also, these PROTACs fail to dock into the pocket of the CRBN-BRD4 complex.

DiffLinker takes 3D fragments as input, and in order to compare with our model, pocket parameters are not considered in this trial. The number of heavy atoms of the linker can be manipulated, while the anchor cannot be predefined, regardless of the anchor position parameter (Github commit number: d4783f5). Therefore, the program determines the anchor positions with the shortest distance, resulting in false spatial orientations. The erroneous linker structures will cause the designed PROTAC molecules unable to dock and unable to form the correct ternary complex.

### Docking Validation

The CRBN-BRD4 protein and 524 postprocessed molecules are input data for docking validation, and all these input data are presented on our GitHub. Molecular docking was performed by the AutoDock4[6] program using the implemented empirical free energy function and the Lamarckian Genetic Algorithm (LGA).[7] In general, the docking parameters for

Table S2: Performance of generating unique and novel bridging molecules in comparison to other ML methods. The preparation and filter steps are identical to AIMLinker.

| Filters | AIMLinker | DeLinker | 3DLinker | DiffLinker |
|---|---|---|---|---|
| Number of outputs from network | 2,000 | 2,000 | N/A | 2,000 |
| Accurate anchor positions | 2,000 | 2,000 | N/A | 0 |
| Number of outputs after removing duplications | **1,728** | 1,499 | N/A | N/A |
| Numer of outputs after removing unlinked structures | **1,175** | 883 | N/A | N/A |

AutoDock4 were kept to their default values. The docking grid was defined by the PROTAC binding site of the CRBN-dBET6-BRD4 (PDB: 6BOY).[8] All the rotatable bonds in ligands are flexible during the docking procedure, and we kept all the protein residues inside the binding pockets rigid. Based on the docking results, we applied the root-mean-square deviation (RMSD)[9] algorithm on each result and arranged the top 10 best RMSD results in a SD file on our GitHub.

### Development Environments

AIMLinker is mainly integrated with Python 3.6 with main packages including TensorFlow-GPU 1.15, rdkit 2022.03.3, joblib 0.13.2, and matplotlib 3.3.4. During docking validation, AutoDockTools 1.5.7 was used to determine the polar hydrogens, the Gasteiger charges and the docking grid, and AutoDock 4.2.6 was applied as a docking algorithm. For ligands and proteins visualization, we use Ketcher 2.5.0 and DSV 21.1.0.20298, while the latter also plays an important role in data preparation.
